# Divergence of trafficking and polarization mechanisms for PIN auxin transporters during land plant evolution

**DOI:** 10.1101/2022.04.28.489888

**Authors:** Han Tang, Kuan-Ju Lu, YuZhou Zhang, You-Liang Cheng, Shih-Long Tu, Jiří Friml

## Abstract

The phytohormone auxin and its directional transport through tissues play a fundamental role in development of higher plants. This polar auxin transport predominantly relies on PIN-FORMED (PIN) auxin exporters. Hence, PIN polarization is crucial for development, but its evolution during the rise of morphological complexity in land plants remains unclear. Here, we performed a cross-species investigation by observing the trafficking and localization of endogenous and xenologous PINs in two bryophytes, *Physcomitrium patens* and *Marchantia polymorpha*, and in the flowering plant *Arabidopsis thaliana*. We confirmed that the GFP fusion did not compromise the auxin export function of all examined PINs by using radioactive auxin export assay and by observing the phenotypic changes in transgenic bryophytes. Endogenous PINs polarize to filamentous apices, while xenologous Arabidopsis PINs distribute symmetrically on the membrane in both bryophytes. In Arabidopsis root epidermis, bryophytic PINs show no defined polarity. Pharmacological interference revealed a strong cytoskeleton dependence of bryophytic but not Arabidopsis PIN polarization. The divergence of PIN polarization and trafficking is also observed within the bryophyte clade and between tissues in individual species. These results collectively reveal a divergence of PIN trafficking and polarity mechanisms throughout land plant evolution and a co-evolution of PIN sequence-based and cell-based polarity mechanisms.

## Introduction

Auxin is a crucial regulator of polarity and morphogenesis in land plants (Kato et al., 2018; Leyser, 2018; Mockaitis and Estelle, 2008; Smit and Weijers, 2015; Yu et al., 2022). The auxin gradients and local maxima within tissues coordinate a broad spectrum of plant development, ranging from embryogenesis to organ formation and tropisms (Friml, 2021; Vanneste and Friml, 2009). The establishment of the auxin gradient relies dominantly on directional auxin transport driven by the efflux carriers, PIN-FORMEDs (PINs). In different tissues, specific PINs are polarized at different plasma membrane (PM) domains, thus directly driving the directionality of auxin flow (Adamowski and Friml, 2015). Therefore, given the essential impact of auxin flow in various developmental processes, the function and polarization of PIN proteins are crucial for maintaining the correct pattern of plant growth and patterning (Sauer and Kleine-Vehn, 2019).

*PINs* are found in all land plants and can be traced back to charophytic green algae (Skokan et al., 2019; Viaene et al., 2013). The functional conservation of PINs in the auxin transport has also been demonstrated by exogenously expressing PINs from the green algae *Klebsormidium flaccidum (K. flaccidum*), the moss *Physcomitrium patens (P. patens*), and the angiosperm *Arabidopsis thaliana* in transgenic plants and in heterologous systems (Skokan et al., 2019; Zourelidou et al., 2014). Additionally, when xenologous PINs from charophytes or Arabidopsis are overexpressed in *P. patens*, the transgenic plants show growth inhibition that resembles auxin deprivation phenotypes (Lavy et al., 2016; Skokan et al., 2019; Tao and Estelle, 2018; Viaene et al., 2014). Therefore, these observations support the idea that PIN-mediated polar auxin transport may govern plant development since the emergence of land plants.

PIN polarity regulations have been extensively investigated in the angiosperm Arabidopsis. Canonical PINs, featuring a long central hydrophilic loop (HL) between two transmembrane domains, are delivered via the ER-Golgi vesicle trafficking pathway to the plasma membrane (PM). De-polymerization of actin filaments induces the accumulation of PIN-labelled small intracellular puncta near the PM, but with no apparent PIN polarity defect (Geldner et al., 2001; Glanc et al., 2018; 2019). Microtubules are involved in the cytokinetic trafficking of PINs, but are not required for the polarity establishment or maintenance at the PM of non-dividing cells (Geldner et al., 2001; Glanc et al., 2019; Kleine-Vehn et al., 2008b). Notably, the disruption of both cytoskeletal networks delays but does not abolish AtPIN2 polarization in Arabidopsis epidermal cells (Glanc et al., 2019). This suggests that the cytoskeletal networks are involved, but not strictly essential for PIN polar trafficking, while other mechanisms contribute to the PIN polar localization. PINs are known to undergo constitutive cycles of endocytosis and recycling, which is modulated by auxin itself (Narasimhan et al., 2021; Narasimhan et al., 2020); this process is essential for their polar distribution (Doyle et al., 2015; Kleine-Vehn et al., 2011).

The Phosphorylation of specific sites within the HL region has been demonstrated to be a critical determinant for PINs’ apical-basal polarization in Arabidopsis epidermis. A serine/threonine kinase, PINOID (PID), phosphorylates specific sites on AtPIN2 and leads to its apical localization (Friml et al., 2004). In contrast, when phosphatase 2A (PP2A) de-phosphorylates AtPIN2, which counteracts the PID-dependent phosphorylation, and thus guides the delivery of AtPIN2 to the basal domain of epidermal cells (Michniewicz et al., 2007). The phosphorylation sites targeted by different kinase families are crucial for polar localizations and functions of PINs, and most sites are highly conserved within Arabidopsis canonical PINs (Zwiewka et al., 2019). Since PINs are present in all land plants, one can hypothesize that the phosphorylation-based polarity regulation may have been established since the emergence of early land plants. However, it has never been clarified whether these phosphorylation sites are evolutionary conserved in early land plants.

The PIN polarization has been observed in the early-divergent moss, *P. patens*, which grows as filamentous protonemata, and its endogenous PpPINA-GFP shows a polar localization at the tip of protonema cells (Bennett et al., 2014; Viaene et al., 2014). However, the polar localization of PpPINA-GFP is not robust in other species. When PpPINA-GFP is expressed in Arabidopsis root epidermal cells, where AtPIN2-GFP presents a clear apical localization, PpPINA-GFP is mislocalized at both basal and apical sites (Zhang et al., 2019). Furthermore, PINs from the liverwort, *Marchantia polymorpha (M. polymorpha*), and from the green algae, *K. flaccidum*, are also mislocalized in root epidermal cells of Arabidopsis (Zhang et al., 2019). These distinctive PIN localizations in different species suggest that mechanisms for PINs trafficking and polarization may have diversified after the emergence of land plants. Despite the profound significance of PIN polarization and the resulting directional auxin transport for land plant development, PIN trafficking and polarization mechanisms are mainly derived from the angiosperm model *Arabidopsis thaliana*.

Here we investigated PIN trafficking/polarization mechanisms from an evolutionary perspective. We show that canonical PINs from bryophytes *P. patens, M. polymorpha*, and Arabidopsis present high conservation in their transmembrane domains and phosphorylation sites. Endogenous PIN-GFPs show different localization patterns in various developmental contexts, suggesting tissue-specific PIN polarization mechanisms. Cross-species investigation revealed that xenologous PINs can traffic to the PM but fail to enrich at the polar domains revealing species-specific mechanisms for PIN polarization. This notion was further verified by different dependency of cytoskeletons for the polarization of Arabidopsis PINs and bryophytic PINs. Overall, our results highlight that PINs trafficking and polarization mechanisms underwent complex evolution during the gradual rise of morphological complexity in land plants.

## Results

### Phosphorylation sites are highly conserved between bryophytic and Arabidopsis PINs

To dissect the conservation between bryophytic and Arabidopsis PINs, we first performed a phylogenetic analysis. The coding sequence of the single canonical PIN, MpPINZ, from *M. polymorpha*, three canonical PINs, PpPINA-PpPINC, from *P. patens*, and five canonical PINs, AtPIN1-AtPIN4 and AtPIN7, from Arabidopsis were subjected to MEGA X program (Kumar et al., 2018) for the phylogenetic study. Bryophytic PINs are categorized in one group, and the MpPINZ is closer to Arabidopsis PINs than PpPINs are (Figure 1A). Arabidopsis AtPIN1 and AtPIN2 were identified with clear polarity at the PM that plays crucial role in embryogenesis, organ formation, and tropic growth (Krecek et al., 2009; Omelyanchuk et al., 2016). The phylogenetic tree showed that both AtPIN1 and AtPIN2 are equally close to bryophytic PINs, while AtPIN2 was extensively investigated for its polarization in roots (Abas et al., 2006; Glanc et al., 2018; Kleine-Vehn et al., 2008a), thus we first picked AtPIN2 as our reference for further alignment analyses. The identity index of coding amino acid sequences showed that full-length AtPIN2 has around 50% identity with each bryophytic PIN (Figure 1B). Since the transmembrane domains show a high similarity between all examined PINs, we suspected that central HLs are more divergent. Surprisingly, the identity index for only the HL region of AtPIN2 presented over 40% identity with MpPINZ (45%), PpPINA (42%), and PpPINB (43%) (Figure 1B). The identity index of full-length AtPIN1 and HLs showed similar results as AtPIN2 (Supplemental Figure 1).

**Figure 1.**
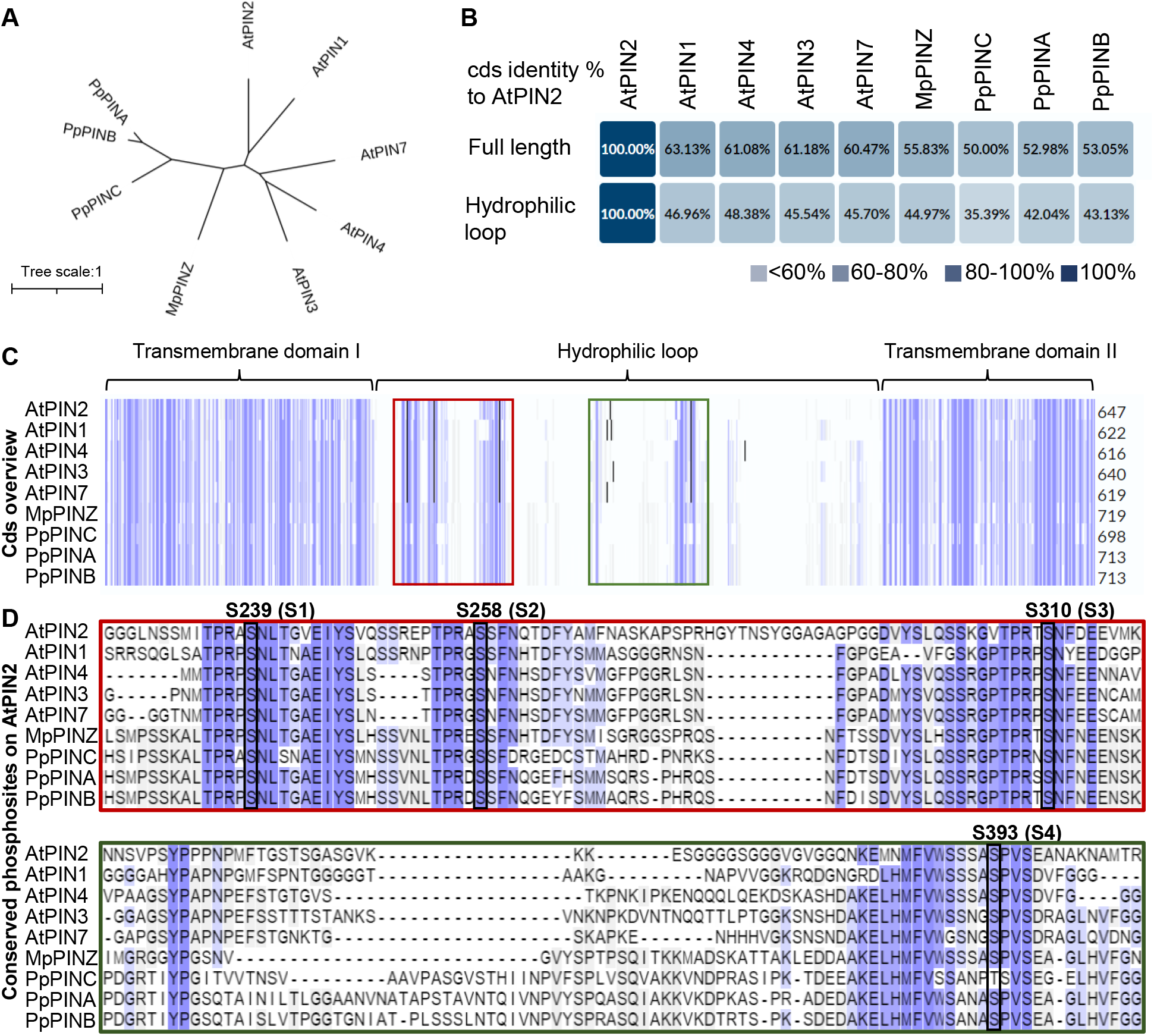
Phosphorylation sites are highly conserved between bryophytic and Arabidopsis PINs. (A) Phylogenetic analysis of canonical PINs from early-divergent plants *P. patens (Pp), M. polymorpha* (*Mp*), and representative angiosperm *A. thaliana* (*At*). The unrooted tree shows the relationships of different PINs in two representative bryophytes and Arabidopsis. The scale bar represents the number of changes per site. (B) Identity indexes of all PINs compared to AtPIN2 with full length or only the hydrophilic loop region of coding amino acid sequences. Identity indexes of all PINs compared to AtPIN1 is shown in Supplemental Figure 1. (C) Alignment of PIN amino acid sequences. Identical amino acids are highlighted with blue column. The rea and green boxes show the hydrophilic loop regions bearing conserved phosphorylation sites, which are enlarged in (D). (D) Four conserved phosphorylation sites, labeled by S1 to S4, were verified in previous AtPIN2 studies and are depicted with black frames. Note that S1 to S4 are conserved in every examined PINs.

The overview of coding sequences for all examined PINs revealed highly conserved transmembrane domains at N-terminus and C-terminus, connected by a less conserved HL region (Figure 1C). Because the polarization of AtPIN2 is tightly associated with its phosphorylation status on the HL, we used AtPIN2 as a reference to search for and highlight these experimental-identified phosphorylation sites (Barbosa et al., 2018; Sukumar et al., 2009; Zhang et al., 2010). Comparing to transmembrane domains, despite of a relatively lower conservation in their HLs, the four identified phosphorylation sites are fully conserved between Arabidopsis PINs and bryophytic PINs (Figure 1D), which suggests that the PIN phosphorylation might be evolutionarily conserved for the regulation of their intracellular localization.

### The hydrophilic loops in AtPIN2, PpPINA, and MpPINZ are less conserved

To dissect how conserved between Arabidopsis PINs and bryophytic PINs is in their protein structures, we next performed structural prediction and alignment. Structures predicted by the Alphafold2 program resemble newly resolved structures of AtPIN1 and AtPIN8 (Supplemental Figure 2) (Jumper et al., 2021; Ung et al., 2022; Yang et al., 2022), supporting that the structure prediction is reliable. Therefore, to compare the structures of well-investigated polarized AtPINs, we predicted the structures of AtPIN1 and AtPIN2 by Alphafold2 and aligned their predicted structures in ChimeraX (Jumper et al., 2021; Pettersen et al., 2021). The structural conservation was presented in almost perfect alignment of transmembrane domains while their HL regions showed similar patterns (Figure 2A, close arrowheads) with two extra random loops in AtPIN2 (Figure 2A, open arrowheads). We next aligned the structure of AtPIN2 with PpPINA and MpPINZ. The transmembrane domains are highly conserved with nearly perfect alignments, but the HLs are less conserved that only one loop showed partial similarity (Figure 2B, close arrowheads). Note that to reveal the conservation and phosphorylation sites clearly, the orientation of protein structures presented in Figure 2A and 2B-E were rotated in different angles. In general, bryophytic PINs possess loosen and larger loops compared to AtPIN2 (Figure 2B-2E). The predicted structures were shown individually with the annotated phosphorylation sites that are indicated in Figure 1D (Figure 2C-2E). The structural conservations in transmembrane domains imply that ancestry bryophytic PINs may traffic to the PM through the same delivery pathways as Arabidopsis PINs, whereas the loosen loops with conserved phosphorylation sites suggest a deviation from conserved regulation for their polarization.

**Figure 2.**
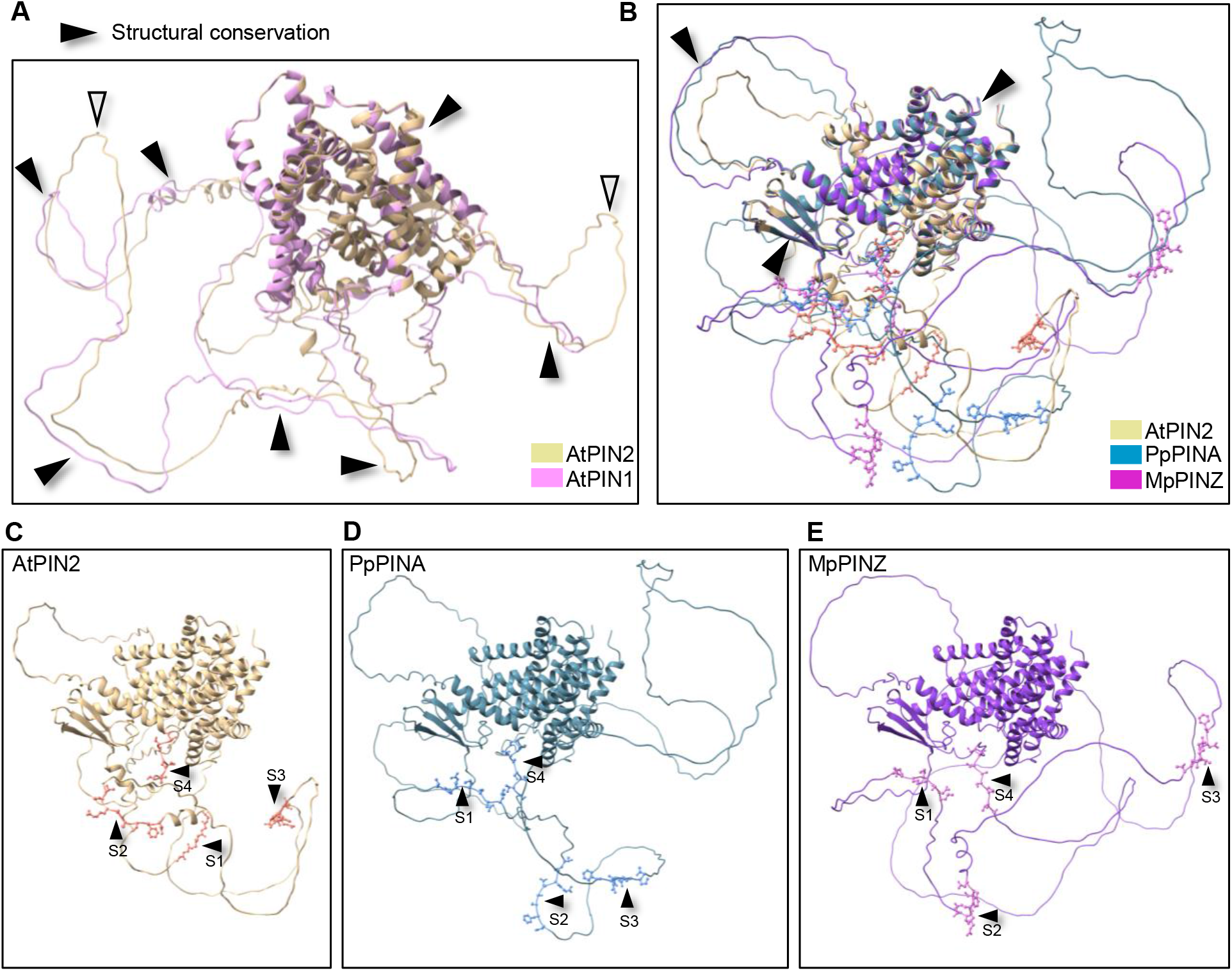
The hydrophilic loops in AtPIN2, PpPINA, and MpPINZ are less conserved. (A) Structural alignment of AtPIN1 and AtPIN2. The protein structures are predicted by Alphafold2 and aligned in ChimeraX software. The structural conserved regions are indicated by black arrowheads, and the non-conserved regions are indicated by empty arrowheads. (B) Structural alignment of AtPIN2 with PpPINA and MpPINZ. Only transmembrane domains and one loop aligned with each other, while the majority of hydrophilic loop regions are not conserved. The four conserved phosphorylation sites are labeled with atom details in ball-and-stick style. (C) – (E) Individual protein structure retrieved from (B). The four conserved phosphorylation sites are indicated by arrowheads.

### GFP-fused PIN proteins possess auxin export activity

The sequence and structural analyses between Arabidopsis PINs and bryophytic PINs revealed high conservation in sequence, phosphorylation sites, and structures of transmembrane domains. Thus, we next would like to investigate whether bryophytic PINs are delivered to the PM with polar domain enrichment as Arabidopsis PINs. To achieve this, we first generated PINs with GFP-fused into an equivalent position of AtPIN1, AtPIN2, PpPINA, and MpPINZ as shown in Figure 3A (Zhang et al., 2019). To verify the auxin export function of these GFP-fused PINs, we subcloned each GFP-fused *PIN* gene into a moss vector, where each *PIN-GFP* gene was driven by an inducible *XVE* promoter (Kubota et al., 2013). We generated transgenic moss plants carrying different *XVE::PIN-GFPs* separately, and we used these transgenic lines to perform the auxin export assay. In brief, the overexpression of *PIN-GFPs* were induced for 3 days, followed by a radioactive auxin H^3^-IAA treatment for 24 hours. The radioactive tissues were then washed twice and were incubated in fresh growth mediums for another 24 hrs. The culture medium was collected for H^3^ scintillation detection (Lewis and Muday, 2009). The wild-type moss plants were taken along as an internal control to show the basal exportation of H^3^-IAA by endogenous PpPINs. In comparison to the wild-type, all examined PIN-GFP plants showed higher amount of radioactive auxin in the culture medium, indicating their auxin export activities (Figure 3B).

**Figure 3.**
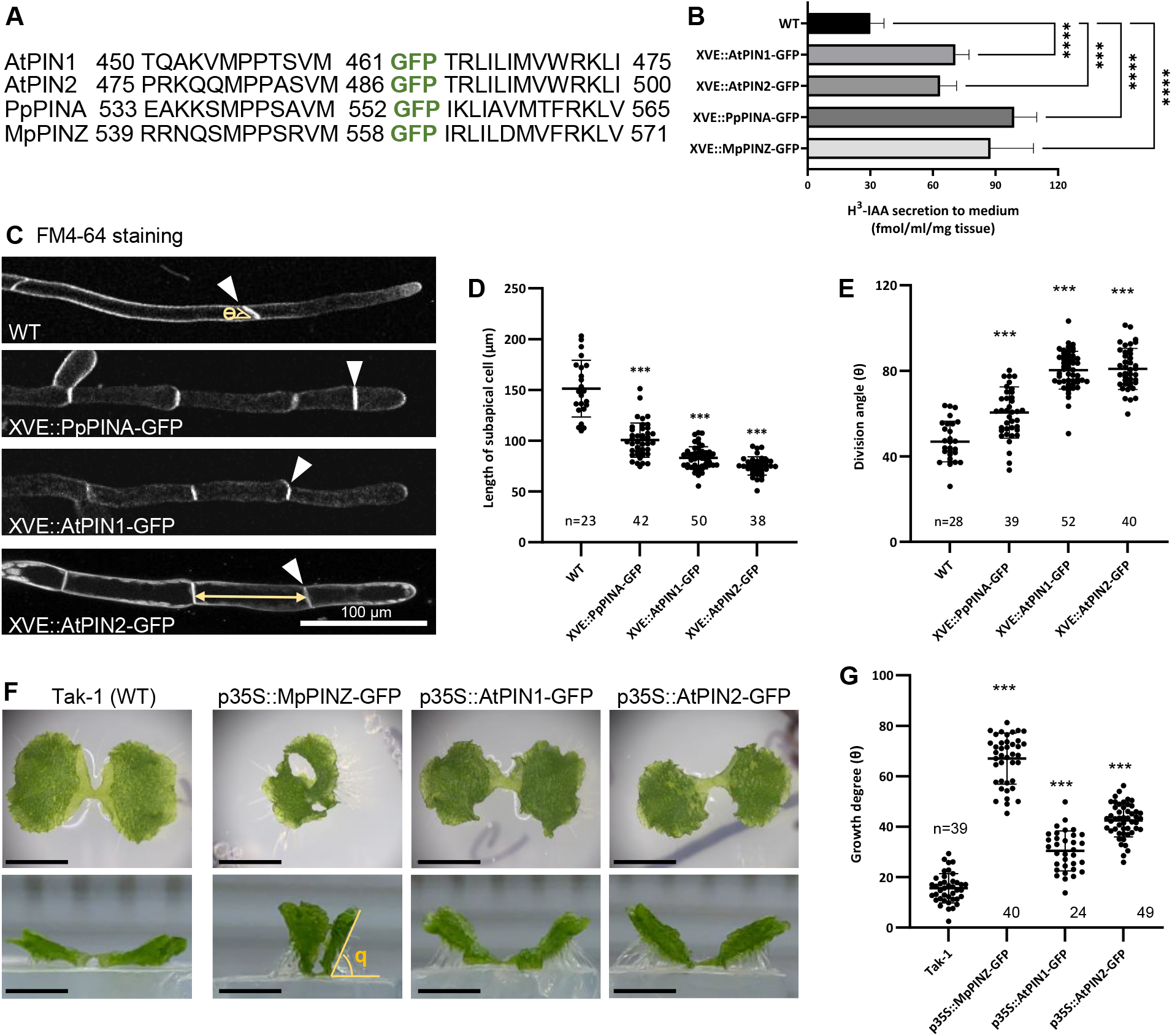
GFP-fused PIN proteins possess auxin export activity. (A) The insertion site of GFP in indicated PIN proteins. Numbers represent the amino acid position in respective proteins. (B) The auxin export assay with *P. patens* transgenic plants. Fresh tissues were incubated with radioactive H^3^ IAA, followed by washing and the radiative H^3^-IAA exported in the new culture medium was measured with the scintillation detector after one day. (C) Representative protonema cells of the indicated genetic background. The cell outline was stained with FM4-64. White arrowheads indicate the first cell division plane, and the yellow double arrow indicates the cell length of the subapical cell. Scale bar = 100 μm. (D) and (E), Quantitative results of the subapical cell length and the division angle (ϴ) of indicated lines, respectively. Bold horizontal lines indicate the median and whiskers indicate the first and the third quarter of the data value. ***P<0.001; Student’s *t*-test. (F) Above (upper panels) and side (lower panels) views of indicated *M. polymorpha* lines. Tak-1 (WT) expands its thallus horizontally on the agar surface. The overexpression of MpPINZ-, AtPIN1-, and AtPIN2-GFP showed vertical thallus growth, which generated a large angle between the lower surface of the thallus and the surface of the agar (ϴ). Scale bar = 0.5 cm. (G) The quantification of the thallus growth angle (ϴ) showed in A. Bold horizontal lines indicate the median and whiskers indicate the first and the third quarter of the data value. ***P<0.001; Student’s *t*-test.

Additionally, the function of PIN-GFPs was also confirmed in the growth changes caused by *PIN-GFPs* overexpression in both *P. patens*. and *M. polymorpha*. During early development, *P. patens* gradually transits its filamentous protonemata from thicker/shorter chloronema cells with perpendicular division planes to thinner/longer caulonema cells with oblique division planes (Rensing et al., 2020). In the previous study, PpPINA, PpPINB, and PpPIND overexpression has been demonstrated to inhibit the protonema transition (Viaene et al., 2014). Here we showed that overexpression of *PpPINA-, AtPIN1-*, and *AtPIN2-GFP* led to similar defects in this chloronema-caulonema transition with a shorter cell length of the subapical cell and a smaller division angle (Figure 3C-3E). The wild-type *M. polymorpha* has prostrate thalli; however, with the overexpression of either *MpPINZ-GFP* or *AtPIN-GFPs*, the thallus grew more vertically, as apparent from the side view (Figure 3F), phenocopying the auxin deficient phenotype (Kato et al., 2017). The angles between the thallus and the horizontal agar were measured to quantify the vertical growth (Figure 3G). The overexpression of *MpPINZ-GFP* caused the most striking phenotype, but the overexpression of *AtPIN-GFPs* also resulted in a vertical growth that showed significant differences from the wild-type. Our results show that all PIN-GFPs can export auxin in the moss system and cause phenotypic changes in both *P. patens* and *M. polymorpha* upon overexpression.

### Endogenous PINs present different localization patterns in different types of tissue

To observe the PpPINA-GFP localization in *P. patens*, we used a stable moss transgenic line carrying the native promoter controlled *PpPINA* genomic DNA-GFP fusion (*pPINA::PpPINA-GFP*) where GFP was inserted in the same position as we verified. The moss *P. patens* has a filamentous protonema stage and a leafy gametophore stage in its life cycle. The initial protonema cell was newly regenerated from a detached leaf, and the elongated protonema was imaged from a 6-days-old moss colony. PpPINA-GFP localized at the PM of the protonema tip with a clear polarity in both initial and elongated protonemata (Figure 4A). To examine whether the polar localization of PpPINA-GFP appears in complex tissues composed of multiple cell layers as well, we observed its localization in gametophytic leaves. In young gametophytic leaves, PpPINA-GFP showed clear basal-apical polarization along the leaf axis with notable corner enrichment (Figure 4A), whereas in mature leaves PpPINA-GFP was evenly distributed on the PM (Supplemental Figure 3). We also analyzed another endogenous *P. patens* PpPINB, fused with GFP and driven by its endogenous promoter. Notably, PpPINB-GFP did not show visible polarity at the tip of protonema, but more even PM distribution with an increased intracellular signal (Supplemental Figure 4). The difference between the localization of PpPINA-GFP and PpPINB-GFP suggests different trafficking and polarization pathways for different PINs co-existing in the same cell already in bryophytes.

**Figure 4.**
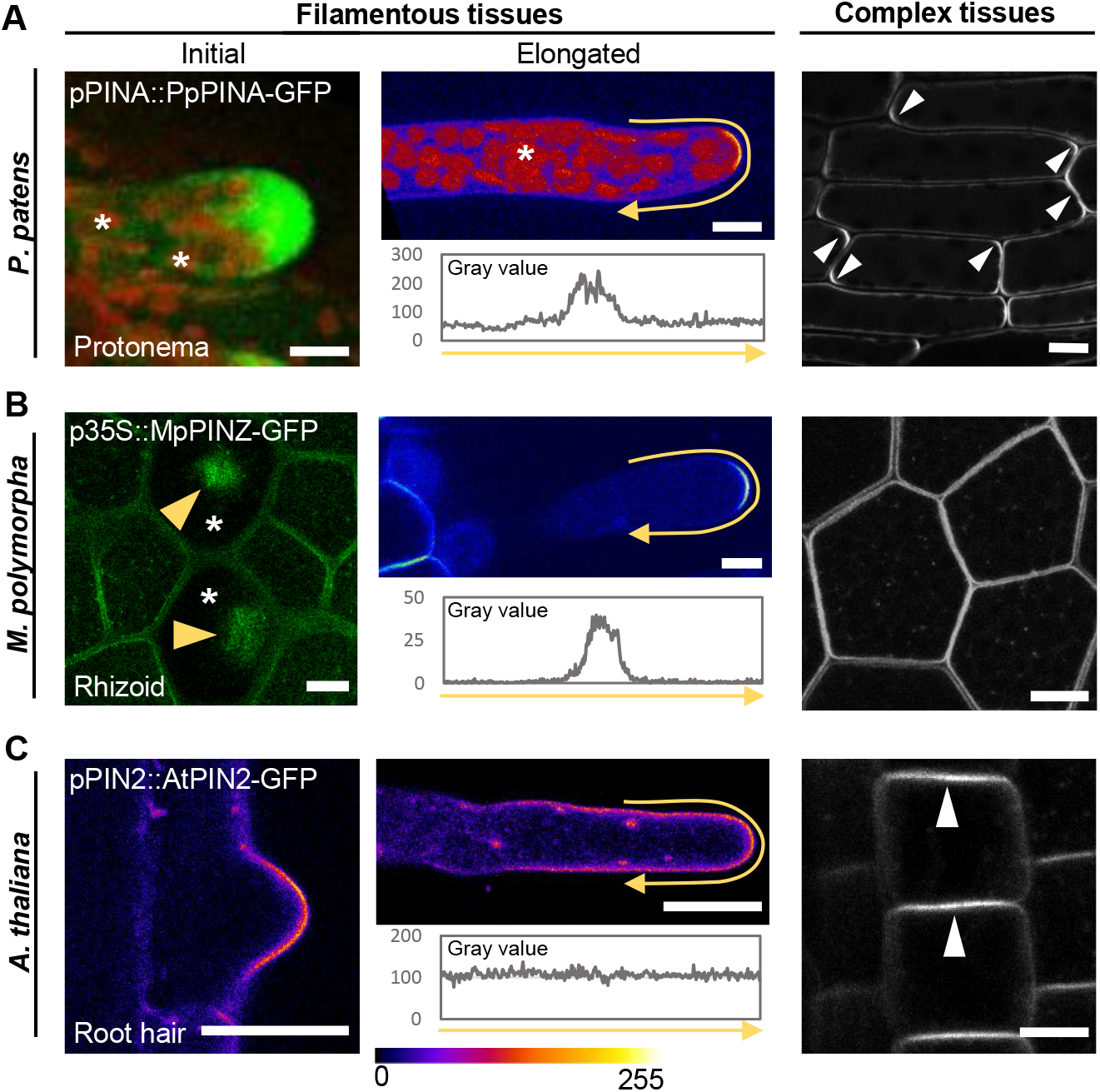
Endogenous PINs present different localization patterns in different tissues. (A) The localization of PpPINA-GFP in tissues with different complexities. PpPINA-GFP is polarized to the tip of initial and elongated protonema cells. The initial protonema cell is regenerated from a detached leaf and the representative image shows the maximum projection with a 5 μm thick Z-section. Auto-fluorescence of the chloroplasts is indicated by asterisks. In elongated protonema cell, the polarity of PpPINA-GFP is plotted by a intensity measurement along the PM as the yellow arrows represented. The same measurement is applied to (B) and (C). In complex tissues composed of multiple cell layers e.g. young leaves in *P. patens*, PpPINA-GFP is polarized at both apical and basal domains (white arrowheads). Scale bars = 10 μm. (B) The localization of MpPINZ-GFP in emerging rhizoids, young rhizoids, and gemma epidermal cells. Representative image with a 5 μm thick Z-section at the gemma surface shows the accumulation of MpPINZ-GFP at tips of emerging rhizoids as indicated by yellow arrowheads in rhizoid precursor cells (asterisks). MpPINZ-GFP is polarized at the tip of young rhizoids (middle). A middle section image showing the even distribution of MpPINZ-GFP on the PM of gemmae composed of multiple cell layers. Scale bars = 10 μm (C) AtPIN2-GFP shows a polarized signal at the tip of the initial root hair but the polarized signal is not observed in elongated root hairs. All imaging details are described in Methods. The pixel values ranging from 0 to 255 are represented by the rainbow color. In Arabidopsis epidermal cells, AtPIN2-GFP shows apical polarization as white arrowheads indicated. Scale bar, 10 μm for root hairs, and 1 cm for epidermis.

The divergence of PpPINA-GFP polarity in filamentous protonema cells and in gametophytic leaves made us wondering if this tissue-specific polarization of PINs is conserved in other bryophytes. To achieve this, we looked at another bryophyte, the liverwort *M. polymorpha* that produces gemmae as the asexual reproductive progenies consisted of multiple cell layers. After water imbibition, single-cellular rhizoids emerge from the large rhizoid precursor cells on epidermis of a gemma (Shimamura, 2016). To analyze the subcellular localization of the sole canonical PIN, MpPINZ, in *M. polymorpha*, we generated a *p35S::MpPINZ-GFP* transgenic line. Interestingly, MpPINZ-GFP localized on the PM with small intracellular puncta and with no apparent polarity in all gemma epidermal cells (Figure 4B, right panel). However, when gemmae were stimulated to grow rhizoids, the signal of MpPINZ-GFP accumulated at the protrusion site of emerging rhizoids in the rhizoid precursor cells (Figure 4B, yellow arrowheads). Later after their emergence, MpPINZ-GFP was polarized at the tip of young rhizoids (Figure 4B), whereas the signal diminished when the rhizoids elongated.

The localization of PpPINA- and MpPINZ-GFP in filamentous and complex tissues suggests distinct polarity recognition mechanisms for bryophytic PINs in different tissue types. To examine if Arabidopsis PINs show similar polar patterns as bryophytic PINs in different types of tissues, we observed *pPIN2::AtPIN2-GFP* line in filamentous root hairs and complex epidermal cells. As previously described, AtPIN2-GFP showed apical localization in epidermal cells (Figure 4C) (Zhang et al., 2019). Since the polarity of bryophytic PINs in their original species mainly appeared at the tip of filamentous tissues, we focused on the localization of AtPIN2-GFP in initial and elongated root hairs, respectively. AtPIN2-GFP revealed a polar localization at the tip of initial root hairs (Figure 4C). However, the polarity of AtPIN2-GFP signals diminished in the elongated hairs (Figure 4C). The different polarity patterns of PINs in different types of tissue support the notion that regulations of PIN polarization are specialized in different cellular profiles and in different developmental contexts.

### Xenologous PINs are PM localized with no defined polarity

Since bryophytic and Arabidopsis PINs demonstrated tissue- and development-specific polarity regulations, we wonder if the conserved phosphorylation sites in the HL are sufficient to drive the polarization of xenologous PINs in other species. To achieve this, we observed the same moss *XVE::PIN-GFPs* transgenic lines we generated for the auxin export assay, carrying PpPINA-, AtPIN1-, and AtPIN2-GFP, respectively. In protonemata, AtPIN1-GFP showed an even distribution on the PM and cell division plane without apparent polarity at the tips, and displayed numerous small intracellular puncta, whereas the PpPINA-GFP signal presented the same polarization as driven by its endogenous promoter (Figure 4A, 5A, and 5D). AtPIN2-GFP under the same induction condition showed much lower intensity but resembled AtPIN1-GFP localization patterns, we thus took AtPIN1-GFP as representative images (Supplemental Figure 5). A short-term induction of *XVE::AtPIN1-GFP* was performed, and it showed no difference in the localization, which verified that the localization patterns of AtPIN1-GFP were not caused by overexpression (Supplemental Video 1).

**Figure 5.**
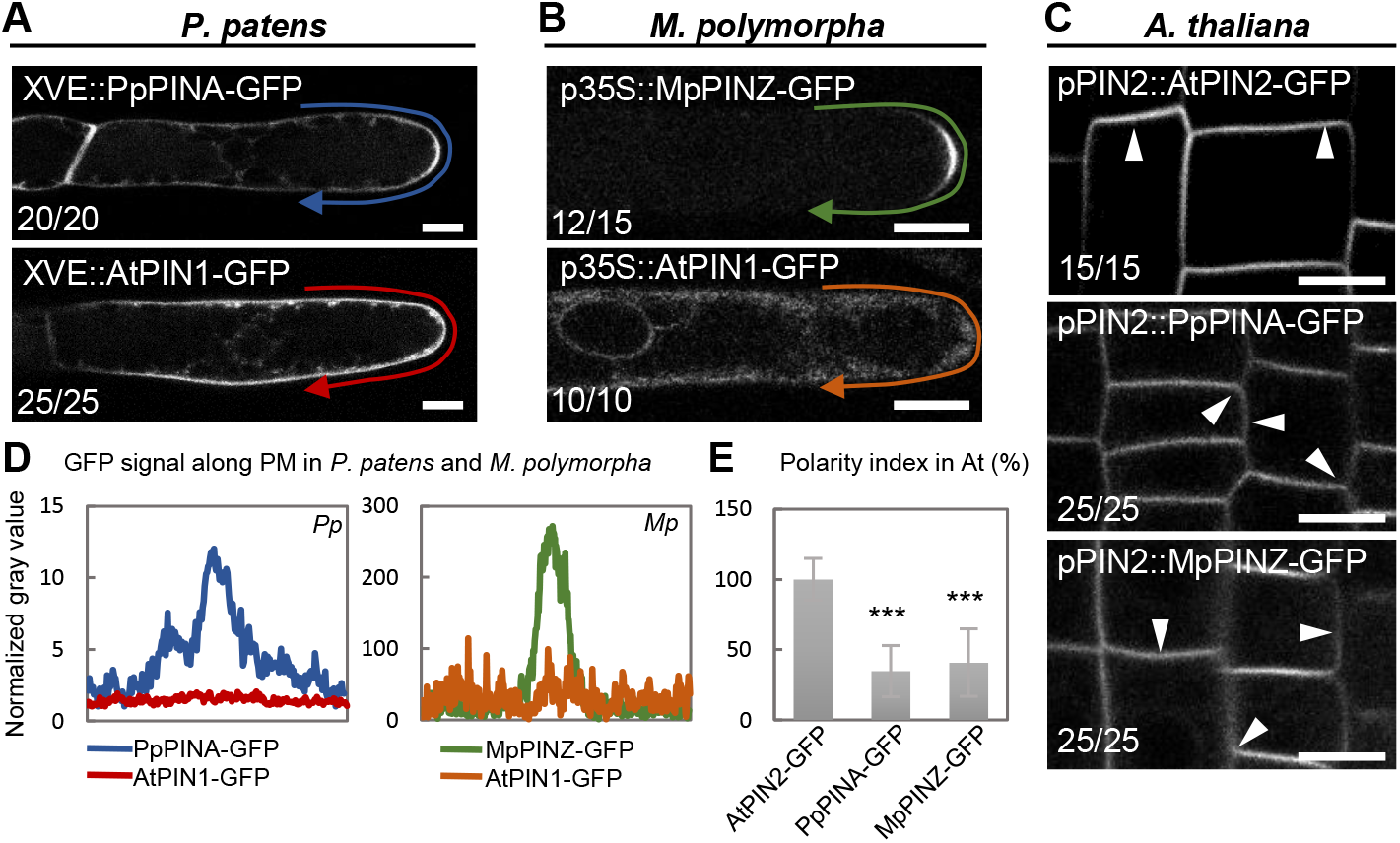
Xenologous PINs are PM localized with no defined polarity. (A) Overexpressed PpPINA-GFP is polarized to the tip of protonema, but overexpressed AtPIN1-GFP is evenly distributed on the PM with intracellular puncta. The occurrence frequency is indicated in the left lower corner in each image. Scale bars = 10 μm. (B) Overexpressed MpPINZ-GFP is polarized to the tip of young rhizoids, while overexpressed AtPIN1-GFP shows strong cytosolic signals and weak PM localization with no polarity. Scale bars = 10 μm. (C) AtPIN2-GFP under its native promoter is apicalized in epidermal cells, but PpPINA-GFP and MpPINZ-GFP are mislocalized to the basal and lateral sites as white arrowheads indicated. Scale bar = 1 cm. (D) The intensity plots of the representative images for PpPINA-GFP, AtPIN1-GFP in *P. patens* protonema cells and MpPINZ-GFP and AtPIN1-GFP in *M. polymorpha* young rhizoids. (E) Polarity index (ratio of signal intensity at the apical plasma membrane/ signal intensity at the lateral plasma membrane) for apicalization of the indicated PIN-GFP in Arabidopsis epidermal cells. PpPINA-GFP and MpPINZ-GFP are significantly lower in the polarity index. 12-20 cells from 3 roots in three independent experiments for each line. ***P<0.001; Student’s *t*-test.

To investigate whether these features are conserved in bryophytes, we introduced AtPIN1-GFP driven by the *35S* promoter into *M. polymorpha*. In gemma epidermal cells, the AtPIN1-GFP showed a non-polar PM localization with high cytosolic signals (Supplemental Figure 6A). In emerging rhizoids, AtPIN1-GFP did not show visible polar localization in rhizoid precursor cells (Supplemental Figure 6B). In young rhizoids, where MpPINZ-GFP showed clear tip polarization, AtPIN1-GFP was evenly distributed on the PM with homogenized cytosolic signals (Figure 5B and 5D).

Since our results indicated that Arabidopsis AtPIN1-GFP in examined bryophytes are delivered to the PM with no visible polarity, we next addressed whether Arabidopsis trafficking machinery can polarize bryophytic PINs. To analyze bryophytic PINs, we observed the protein localization at epidermal cells of transgenic Arabidopsis lines expressing *pPIN2::AtPIN2-, PpPINA-, and MpPINZ-GFP* (Zhang et al., 2019). As previously described, AtPIN2-GFP showed apical localization in the epidermal cells, while PpPINA-GFP localized at both the apical and basal sides and MpPINZ-GFP mainly localized at the basal side with some lateral residence. (Figure 5C and 5E) (Zhang et al., 2019). These data thus support that the machinery driving PIN proteins to the PM through a generally conserved cellular trafficking pathway, whereas the PIN polarization is specialized in different species.

### Cytoskeletal networks are important for the polarization of bryophytic PINs

The diversification of PIN polarities in different plant species and tissues suggests that plant cells might utilize various machineries to deliver and maintain PIN proteins to the target side on the PM. To verify this, we examined the necessity of the cytoskeletal networks in the polarization of bryophytic PINs and AtPIN2 in their original species. We depolymerized actin filaments or microtubules by treating plant tissues with Latrunculin B (LatB) or Oryzalin (Ory), respectively. In *P. patens*, the abolishment of actin filaments resulted in the hyper-polarization of PpPINA-GFP, where PpPINA-GFP accumulated at a focal locus to the very tip of the cell (Figure 6A and 6C). The disruption of microtubules resulted in less accumulation of PpPINA-GFP at the tip of protonemata, and the PpPINA-GFP appeared to be detached from the PM (Figure 6A and 6C). The changes of PpPINA-GFP localization in response to drug treatments demonstrated the requirement of cytoskeletal networks for PpPINA-GFP polarization in filamentous tissues.

**Figure 6.**
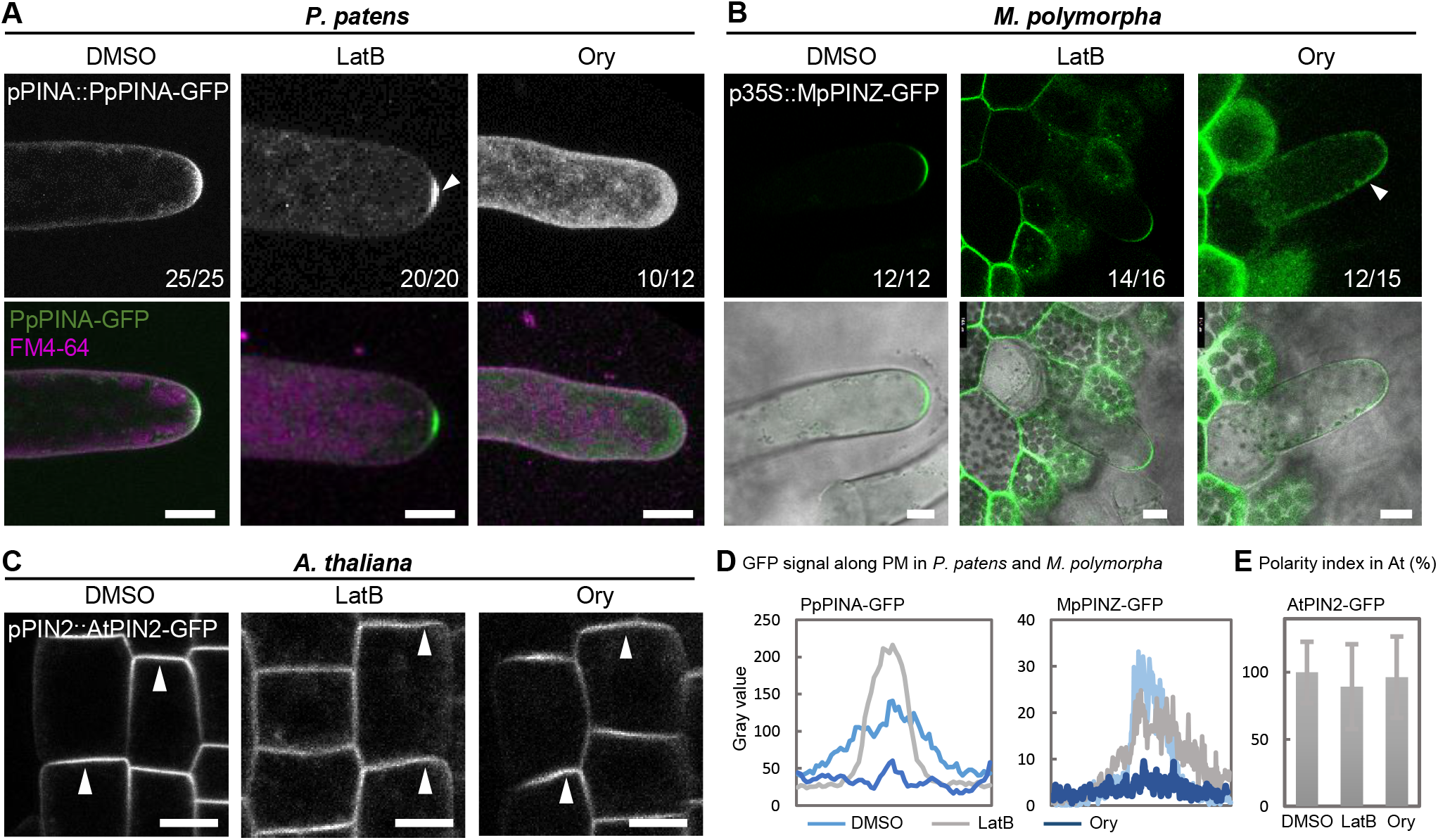
Cytoskeletal networks are important for the polarization of mainly bryophytic PINs. (A) PpPINA-GFP is polarized to the tip of protonema, and the disruption of actin filaments by Latrunculin (LatB) caused its hyperpolarization (white arrowhead). The disruption of microtubules by Oryzalin (Ory) disturbed the polarization of PpPINA-GFP. Scale bars = 10 μm. (B) MpPINZ-GFP is polarized at the tip of young rhizoids, and its polarization was only abolished by Oryzalin treatment (white arrowhead). Scale bars = 10 μm. (C) AtPIN2-GFP is polarized at the apical site of epidermal cells, and disruption of either actin filaments or microtubules did not change its polarization (white arrowheads). Scale bars = 1 cm. (D) Relative intensity of PpPINA-GFP and MpPINZ-GFP along the cell membrane with indicated treatment. (E) Polarity index of AtPIN2-GFP in indicated treatment, no significance is shown. The concentration and duration of treatments for each species are described as in Methods.

In *M. polymorpha*, we used the same pharmacological interference to investigate whether the MpPINZ-GFP polarization in young rhizoids relies on the cytoskeletal networks. MpPINZ-GFP remains polarized at the tip when actin filaments were disrupted, while the disruption of microtubules resulted in the dislocation of the polarized MpPINZ-GFP (Figure 6B and 6C). These results collectively demonstrate the diversification on cytoskeletal dependency for PINs polarization within the bryophyte clade.

In Arabidopsis, the disruption of actin filaments by LatB resulted in an increasing appearance of AtPIN2-GFP signals in the cell interior, while the Ory treatment caused a minor effect on AtPIN2-GFP localization (Figure 6D). Upon cytoskeleton disruption, the apical polarization of AtPIN2-GFP in epidermal cells was still evident; suggesting that in contrast to the dependency of cytoskeletal networks for the bryophytic PINs, the polarization of Arabidopsis PINs mainly relies on other trafficking or polarity retention mechanisms. Again, these results reinforce the notion that mechanisms underlying PIN polarization have been expanded and specifically diversified in different species.

## Discussion

### Evolution of sequence-specific determinants of PIN polarity

In flowering plants, as exemplified by Arabidopsis, the five PM-localized canonical PINs show different polarities in different developmental and tissue contexts, thus mediating directional auxin fluxes and generating asymmetric auxin distribution for plethora of developmental processes ultimately shaping the plant form. Originating from a single PIN auxin transporter, such as found in simple filamentous streptophyte algae (Skokan et al., 2019), PIN family radiated during evolution into PINs showing different expression and localization patterns and mediating many developmental and physiological functions.

Ectopic co-expression of PINs in the same cell type such as AtPIN1 and AtPIN2 in root epidermis can lead to diverse polarity patterns (AtPIN1 – basal; AtPIN2 – apical), thus demonstrating simultaneous presence of multiple polarity mechanisms in those cells (Wisniewska et al., 2006). In this study, we showed that in bryophytes, as exemplified by the moss *P. patens*, PpPINA and PpPINB endogenously expressed in protonemata filaments, present different localization patterns (Figure 4 and Supplemental Figure 4). PpPINA shows tip-focused PM localization, whereas PpPINB can be found more spread on the PM and also intracellularly. Whereas, the functional importance of this difference remains unclear, it clearly shows that already in bryophytes, parallel polarity/trafficking mechanisms exists in the same cells, to which different PINs can be recruited, presumably based on specific sequence-based signals.

Notably, the sequence alignment between the HL regions of PpPINA and PpPINB showed 90.31% identity, suggesting that these signals are presumably encoded within the 10% difference regions. Further analysis of the differences between PpPINA and PpPINB or AtPIN1 and AtPIN2 would help to identify the sequence signals for PIN polarity regulations. Despite the identity of the sequence-based signals remain unclear, our observations show that cellular polarity mechanisms and PIN sequence-based polarity signals, which are crucial for diverse developmental roles in flowering plants, started to diversify already in bryophytes.

### Context-specific determinants of PIN polarity

It is well known from Arabidopsis that the same PINs show different localization patterns in different contexts (Vieten et al., 2005). For example, AtPIN2 is apicalized in epidermal cells, while it is localized on the basal side of young cortex cells (Kleine-Vehn *et al*., 2008a). The observations imply that different cell types possess specific trafficking pathways for the same PIN protein. In line with this, our results show that in filamentous cells, e.g., protonemata in *P. patens*, rhizoids in *M. polymorpha*, and root hairs in Arabidopsis, endogenous PIN proteins are polarized at the tip of apical cells (Figure 4A-4C). Notably, MpPINZ-GFP and AtPIN2-GFP signals were diminished when the rhizoids or root hairs elongated, and the PpPINA-GFP was polarized only in young gametophytic leaves but the pattern dispersed in mature leaves. These data collectively demonstrate that PIN polarity is differentially regulated in different developmental contexts.

In complex tissues with multiple cell layers, unlike in filamentous tissues, the PM-localized MpPINZ-GFP did not show polarity in thalli, while PpPINA-GFP and AtPIN2-GFP presented polar localization at the apical-basal domain of the cells (Figure 4A-4C). This data shows that PIN polarity and trafficking mechanisms have evolved with specific modifications in different tissues and cell types, in both angiosperms and early-diverging land plants. This likely reflects different requirements for directional auxin transport in different developmental contexts, and it implies a co-evolution of PIN sequence-based signals and cell type-specific polar sorting and trafficking mechanisms.

### Diversification of PIN trafficking and polarity mechanisms during land plant evolution

The core mechanisms for auxin biosynthesis, auxin signaling, and PIN-mediated auxin transport are evolutionary conserved across land plants (Blazquez et al., 2020; Kato et al., 2018; Sauer and Kleine-Vehn, 2019). However, bryophytes and vascular plants diverged around 450 million years ago and emerged different types of tissues and organs, it is not clear how conserved the PIN polarity regulation is under such drastic changes during land plant evolution. Our cross-species studies revealed that xenologous PINs, such as Arabidopsis PINs in bryophytes and bryophytic PINs in Arabidopsis, are able to traffic to the PM (Figure 5), which suggests that all canonical PINs can be recognized by the general protein transport machinery in other species.

Interestingly, when the Arabidopsis PINs are ectopically expressed in bryophytes, they fail to form any specific polarity, whereas bryophytic PINs remain in apical-basal domains in Arabidopsis epidermis (Figure 5). These observations may hint an evolutionary loss of some regulatory motifs for PIN polarization that are presence in bryophytic PINs but absence in Arabidopsis PINs. In line with this, the overall coding sequences of bryophytic PINs are longer than Arabidopsis PINs, and the extra sequences are positioned in their HLs, the main regulatory regions for PIN polarization. Our study thus demonstrates that PINs polarity mechanisms are not conserved throughout plant evolution.

This notion was also verified by the differences in cytoskeleton requirements for PIN polarization between bryophytes and angiosperms. The polarity of bryophytic PINs was disrupted when cytoskeletal networks are depolymerized, while Arabidopsis AtPIN2 was not affected (Figure 6). This finding suggests a gradual shift in the dependence of PIN polarity and trafficking from the cytoskeleton-dependent pathways towards the cytoskeleton-independent pathways along the land plant evolution.

Overall, our results demonstrate that different plant species evolved specialized pathways to deliver PINs and maintain their polarity at the PM. This is likely linked to an increasing repertoire of auxin transport developmental roles adopted with increasing morphological complexity during land plant evolution.

## Methods

### Plant growth and transformation

Arabidopsis seeds were surface sterilized and grown on 1/2 MS plates. After a two-day stratification at 4°C, seedlings were grown under the long-day condition (16 hours light, 8 hours dark) at 22°C with 100-120 μmol photons m^−2^s^−1^ white light. For *P. patens*, all transgenic and WT plants were cultured on BCD medium plates in a growth chamber at 24°C under the long-day condition (16 hours light, 8 hours dark) with 35μmol photons m^−2^s^−1^ white light. For *M. polymorpha*, WT and all transgenic plants were cultured on 1/2 B5 plates in a growth chamber under the long-day condition (16 hours light, 8 hours dark) at 22°C with 50-60 μmol photons m^−2^s^−1^ white LED light.

All transgenic plants used in this study are listed in Supplemental Table 1. *P. patens* transgenic plants *pPINA::PINA-GFP* has been generated and verified as previously reported (Viaene et al., 2014), and the inducible overexpression XVE::AtPIN1-GFP, *XVE::AtPIN2-GFP*, and *XVE::MpPINZ-GFP* lines were generated as described in previous studies (Tang et al., 2020; Yamada et al., 2016). In brief, the Gransden 2004 WT moss plants were freshly propagated and transformed via the PEG-mediated transformation (Nishiyama et al., 2000). The transformants were selected under Hygromycin (20μg/mL), and two independent transgenic lines were selected for imaging and analysis.

*p35S::MpPINZ-, AtPIN1-*, and *AtPIN2-GFP M. polymorpha* plants were generated via the Agrobacterium transformation method described before (Kubota et al., 2013). In brief, the apical meristemic region of each two-week-old Takaragaike-1 (Tak-1) thallus was removed and further cut into four pieces. After culturing on 1/2 B5 with 1% sucrose agar plates for 3 days, the cut thalli were transferred to 50ml 0M51C medium with 200 μM acetosyringone (4’-Hydroxy-3’,5’-dimethoxyacetophenone) in 200ml flasks with 130 rpm agitation to coculture with OD600 = 1 density agrobacteria harboring the target construct for another 3 days. The transformed thalli were washed and plated on 1/2 B5 plates with proper antibiotic selection. Independent T1 lines were isolated, and G1 lines from independent T1 lines were generated by sub-cultivating single gemmalings, which emerged asexually from a single initial cell (Shimamura, 2016). The next generation of G1, called the G2 generation, was used for analyses. Arabidopsis transgenic lines bearing bryophytic PIN-GFPs under *AtPIN2* promoter control were generated and used as in a previous study (Zhang et al., 2019). For root imaging, seeds were sown on 1/2 Murashige–Skoog (MS) medium plates and kept at 4°C for 2 days and moved to the growth chamber to culture vertically for another 4 days.

### Plasmid construction

Plasmids and primers for construction and genotype confirmation are listed in Supplemental Table 2. For transgenic lines with inducible overexpression by moss, the insertion site of the *GFP* gene into the HL was indicated in Figure 3A, and the *PIN-GFP* regions were amplified from previously generated plasmids (Zhang et al., 2019) by PCR and cloned into the gateway entry plasmid pENTR/D-TOPO^®^ as the manufacturer suggested. The fragments were sub-cloned into the pPGX8 vector, which contains a p*35S* driven β-estradiol inducible XVE cassette (Floriach-Clark et al., 2021; Kubo et al., 2013; Nakaoka et al., 2012) via a Gateway^®^ LR reaction (Invitrogen) as the manufacture’s recommendation.

To generate a *p35S::MpPINZ-, AtPIN1-*, and *AtPIN2-GFP* construct, the cDNA of the target fragment containing proper *GFP* insertion were amplified with the primers listed in Supplemental Table 2. The amplified fragments were cloned into the pENTR/D-TOPO^®^ vector (Invitrogen) with the protocol supplied by the manufacturer. Plasmids with target genes were further sub-cloned into the pMpGWB102 vector containing a *35S* promoter (Ishizaki et al., 2015) by the Gateway^®^ LR reaction (Invitrogen) as the manufacturer’s recommendation.

### Microscopy

For moss observation, protonemata were cultured in glass-bottom dishes covered by BCD agar medium for 6 – 7d. The live-cell imaging was then performed on a Leica SP8X-SMD confocal microscope equipped with a hybrid single-molecule detector (HyD) and an ultrashort pulsed white-light laser (WLL; 50%; 1 ps at 40 MHz frequency). The Leica Application Suite X was used as a software platform, and imaging was conducted with a HC PL APO CS2 40x/1.20 water immersion objective. The following settings were used: scan speed of 400 Hz, resolution of 1024 × 1024 pixels and standard acquisition mode of the hybrid detector. The time-gating system was activated to avoid the autofluorescence emitted from chloroplasts. For the filament growth assay, the imaging dish was applied with FM4-64 (Invitrogen) solution for 10-30 mins, and the 10x objective lens was used.

For Marchantia observation, gemmae were picked from a gemma cup and transferred into a 24-well plate with 500 μl liquid 1/2 B5 medium. After culturing in the growth chamber for 24 hours, gemmae from each sample were transferred and observed on a slide under Leica Stellaris 8 system with HyD detectors and the ultrashort pulsed white-light laser (WLL; 70%; 1 ps at 40 MHz frequency). The Leica Application Suite X was used as a software platform, and imaging was conducted with an HC PL APO CS2 40x/1.20 water immersion objective. The following settings were used: scan speed of 400 Hz and resolution of 1024 × 1024 pixels. For GFP-containing images, a 488nm white light laser was selected, and the detection range was set between 500nm to 525nm. The tau-gating model, harvesting photons with 1.0-10.0ns lifetime, was used for all Marchantia imaging to avoid autofluorescence emitted from the chloroplasts. For the surface section (rhizoid precursor cell observation), a 5 μm-thick section was set using the z-section method with auto-optimization spacing to capture rhizoids’ protrusion.

For Arabidopsis root imaging, 4-day-old seedlings of each indicated genotype were used for fluorescence imaging. After treatment in the liquid MS medium supplied with indicated chemicals, seedlings were carefully mounted on a slice with growth medium and then placed into a chambered coverslip (Lab-Tek) for imaging. For the root hair imaging, a 3 μm Z-projection image with 1 μm step was taken around the medium plan of the root hair. All fluorescence imaging was performed using a laser scanning confocal microscopy (Zeiss LSM800, 20x air lens). The default setting for GFP detection was applied with 488nm excitation and 495-545nm emission.

### Images quantification

All images were analyzed by Fiji (ImageJ, https://imagej.net/software/fiji/). For the polarization patterns at the tip of filaments in *P. patens*, a line with 5 px thickness was plotted along the PM as depicted in Figure 4 and Figure 5. The representative images for the DMSO control and drug treatments, obtained from the same imaging settings, were used to draw the line, and the mean intensity along the line was shown. For the moss phenotype analysis, mosses bearing inducible *PIN-GFP* were cultivated in the imaging dish for 5 days, followed by 1 μM β-estradiol induction for another 3 days. The cell outlines were stained with FM4-64 for 10-30 mins. The line drawing function in Fiji was used to measure the length of subapical cell, and the line was depicted between the middle points of the two cell division planes. The angle measurement function was applied to examine the angle between the first cell division plane and horizontal cell outline for the division angle measurement.

### Genomic DNA isolation

The genomic DNA of transformants was isolated by the cetyltrimethylammonium bromide (CTAB) gDNA extraction (Schlink and Reski, 2002). In brief, moss tissues were harvested from one full plate and were ground in liquid nitrogen. The ground tissues were then mixed and were incubated with a CTAB buffer, followed by the addition of chloroform. After centrifugation, the supernatant was collected and was precipitated with isopropanol at −20°C for one hour.

### Pharmacological treatments

To depolymerize microtubules and actin filaments, we used oryzalin (Sigma) and latrunculin B (Sigma) to treat plants. The concentration and duration of treatment for different plants have been given in numbers, and the conditions we used have been shown to efficiently depolymerize cytoskeletal networks in different species (Baluska et al., 2001; Baskin et al., 1994; Glanc et al., 2019; Oda et al., 2009; Vidali et al., 2009). For *P. patens*, the chemical was diluted in a liquid BCD medium and was applied to the imaging dishes before imaging.

For *M. polymorpha*, G2 gemmae from transgenic plants were transferred into each well of a 24-well plate containing 500 μl liquid 1/2 B5 medium and cultured in the growth chamber for 16 hours. The chemicals were diluted directly into the medium prior to imaging at the time indicated in the text. Based on previous studies, the concentration of Ory (Era et al., 2013) and LatB (Otani et al., 2018) were selected. For Arabidopsis, 4-days-old seedlings were submerged into the liquid MS medium supplied with chemicals and then transferred to another agar medium for imaging. 0.1% DMSO (Duchefa; 10 mM dimethylsulfoxide) was used as control for all treatments.

### *P.patens* auxin export assay

The auxin export assay performed with transgenic moss plants was modified based on the protocol developed for Arabidopsis seedlings (Lewis and Muday, 2009). In brief, 7-days-old fresh tissues were transfer to liquid BCDAT growth medium with 1μM β-estradiol induction for another 4d with gentle shaking, followed by ^3^H-IAA treatment with a final concentration of 10 nM for 24 hrs. The radioactive tissues were then washed twice with sterile H_2_O, and were cultivated in fresh BCDAT medium for another 24 hrs. The cultivated medium was then collected to mix with ScintiVerse BD cocktail (Fisher, SX18-4) in 1:30 (v:v), and the export of auxin was measured by the Scintillation counter (Beckman, LS6500).

### *M. polymorpha* thallus growth assay

G2 gemmae were transferred onto 1/2 B5 agar plates and grew for 10 days. Gemmae were imaged under a dissecting microscope (SZN71, LiWeng, Taiwan) with a CCD camera (Polychrome M, LiWeng, Taiwan). For measuring the vertical growth angle, an agar cube with an individual plant was cut out from the plate and placed in the middle of a slide. The slide was put on the surface of a laminar flow at a fixed distance to the edge, and images were taken by an HTC U11 cell phone camera. The growth angle was further measured by using ImageJ (https://imagej.net/software/fiji/).

### Phylogenetic analysis

The phylogenetic analysis for full length amino acid sequences of all examined PINs was carried out in MEGA X program (Kumar et al., 2018) and the results were imported into iTOL (https://itol.embl.de/) for visual illustration. The evolutionary history was inferred by using the Maximum Likelihood method and JTT matrix-based model with default settings (Jones et al., 1992). The alignment and identity index were produced by online CLUSTAL alignment program with default settings.

## Author contributions

H. T., K. J. L., and J. F. designed experiments. Y.Z. C. initiated the project and provided key constructs and materials. H. T. performed and analyzed all the *P. patens* and Arabidopsis related experiments with the help of Y. L. C. and S. L. T.. K. J. L. performed and analyzed all the *M. polymorpha* related experiments. H. T., K. J. L, and J. F. wrote the manuscript.

## Acknowledgments

The authors would like to thank Dr. A. Johnson and Ms. Alexandra Mally for their careful proofreading. We also thank Dr. T. Kohchi for sharing the pMpGWB102 vector (Addgene #68556; http://n2t.net/addgene:68556; RRID:Addgene_68556).

