## Supplemental Figures for "Divergence of trafficking and polarization mechanisms for PIN auxin transporters during land plant evolution"

**Figure S1. Identity index of all selected PINs compared to AtPIN1 retrieved from the protein sequence alignment analysis.**

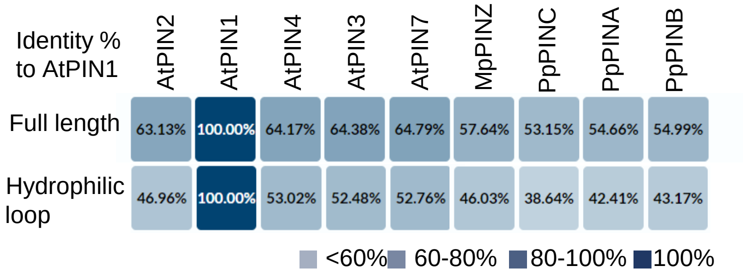


**Figure S1**. Identity index of all selected PINs compared to AtPIN1 retrieved from the protein sequence alignment analysis.

Alignment of full length sequence and hydrophilic loop region for canonical PINs from *Physcomitrium patens* (Pp)*, Marchantia polymorpha* (Mp), and *Arabidopsis thaliana* (At).

#### Figure S2. Structures predicted by the Alphafold2 algorism resemble the newly resolved structures of AtPIN1 and AtPIN8.

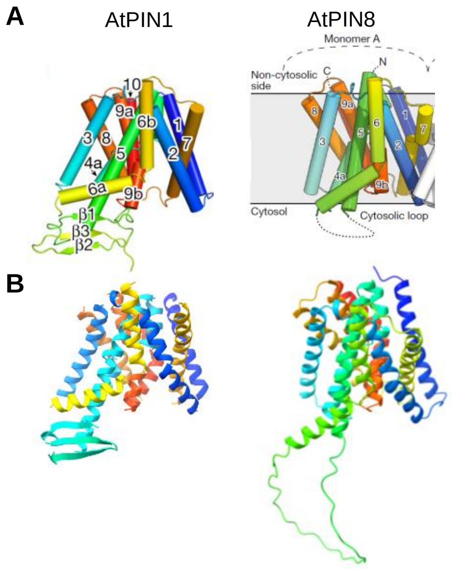


**Figure S2.** Structures predicted by Alphafold2 algorism resemble the newly resolved structures of AtPIN1 and AtPIN8.

(A) Overall structures of AtPIN1 and AtPIN8 monomer modified from Ung *et al*. and Yang *et al*. (Ung *et al.,* 2022; Yang *et al.,* 2022)

(B) Predicted structures of AtPIN1 and AtPIN8 by AlphaFold2.

**Figure S3. PpPINA is evenly distributed on the plasma membrane in mature moss leaves.**

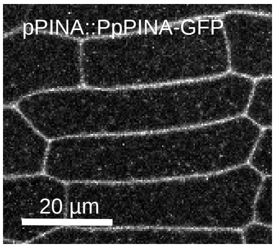


**Figure S3.** PpPINA is evenly distributed on the plasma membrane in mature moss leaves.

As previously reported, PpPINA-GFP showed symmetric distribution in mature gematophytic leaves (Viaene *et al.,* 2014).

**Figure S4. PpPINB is evenly distributed on the plasma membrane with a high cytosolic signal in moss protonema cells.**

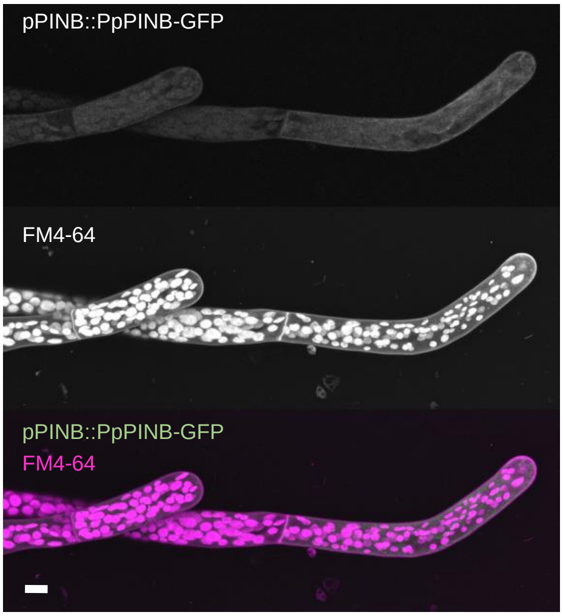


**Figure S4.** PpPINB is evenly distributed on the plasma membrane with a high cytosolic signal in moss protonema cells.

PpPINB-GFP is driven by its endogenous promoter and stained with FM4-64 for cell outline imaging. Scale bar = 10 µm.

**Figure S5. AtPIN2-GFP is localized at the plasma membrane with cytosolic puncta signals in the moss protonema.**

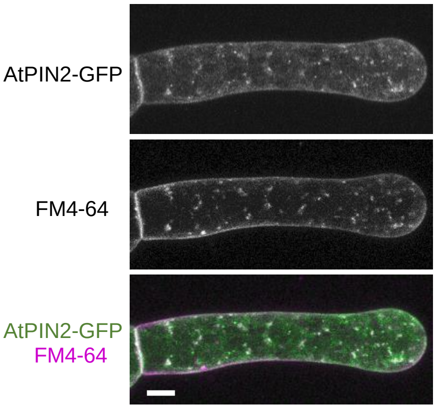


**Figure S5.** AtPIN2-GFP is localized at the plasma membrane with cytosolic puncta signals in the moss protonema.

Moss tissues were cultivated in the imaging dish for 6 days and the expression of AtPIN2-GFP was induced for additional 2 days with 1 µM β-extrodial. The tissues were stained with FM4-64 for 10-30 minutes prior to imaging. Scale bar = 10 µm.

### Figure S6. Arabidopsis AtPIN 1 is not polarized in Marchantia.

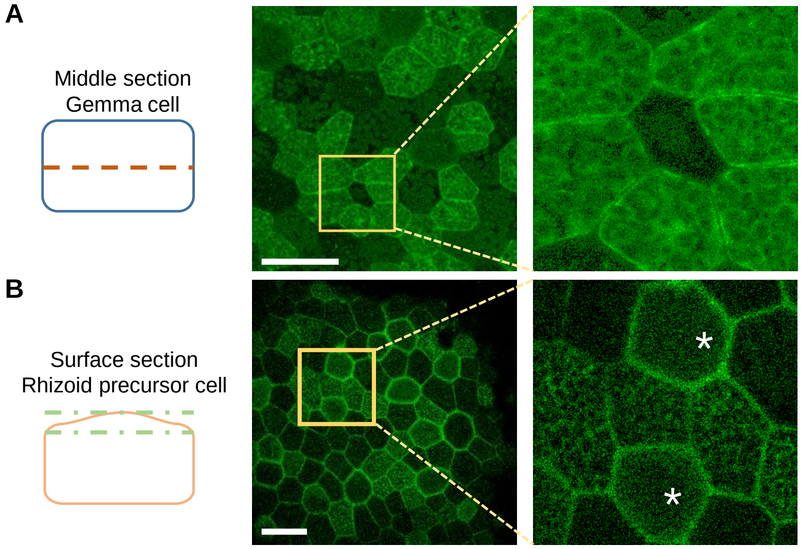


**Figure S6.** Arabidopsis AtPIN 1 is not polarized in Marchantia.

The localization of AtPIN1-GFP in *M. polymerpha* gemma epidermal cells and protrusion site of initial rhizoids (asterisks). Imaging sections were obtained as illustrated. The AtPIN1-GFP is localized at the plasma membrane and in the cytoplasm of all examined tissues with no apparent puncta signal. Scale bar = 50 mm.
